# Synergistic Variants in *C-terminal Binding Protein 1* and *Alkaline Phosphatase* Lead to Mandibular Hypoplasia Through Impaired Wnt Signaling: An Oligogenic Model

**DOI:** 10.64898/2026.09.09.750372

**Authors:** Paul P. R. Iyyanar, Beulah Solivio, K. Nicole Weaver, Rolf W. Stottmann

## Abstract

Craniofacial malformations account for one third of all congenital anomalies. Genetic factors play a vital role, yet the list of causal genes and their mechanisms are far from complete. As part of a larger effort to sequence patients with micrognathia and Pierre-Robin sequence, we identified two candidate pathogenic missense variants in *C-terminal binding protein 1* (*CTBP1*) along with a heterozygous early stop missense variant in *alkaline phosphatase* (*ALPL*) in a proband with mandibular hypoplasia. *Ctbp1* has been shown to regulate Wnt/β-Catenin signaling but it has not yet been implicated in craniofacial development. Here we generated two orthologous variants of *Ctbp1* mimicking the patient variants using genome editing in mice and explored the micrognathia phenotype in combination with a previously reported *Alpl* null allele. *Ctbp1*^*Q148H/G238S*^; *Alpl*^*null/Wt*^ complex heterozygous mutants have smaller mandibles recapitulating the human mandibular hypoplasia. We identified that a reduction in cell proliferation and active β-Catenin levels could possibly account for the micrognathia phenotype in the *Ctbp1*; *Alpl* complex heterozygous. These data uncover a novel role for *Ctbp1* in craniofacial development and highlight the complex genetic and molecular signaling in the pathogenesis of craniofacial malformations.

## Introduction

*C-terminal binding protein* (*Ctbp1*) encodes a multifunctional protein primarily known for its role as a transcriptional corepressor. It plays a vital part in embryonic development, cellular metabolism, and various diseases with multiple reports implicating it in cancer (1). *Ctbp1* is not essential for embryonic survival, but 23% of *Ctbp1* ^*null/null*^ mutants die by postnatal day (P) 20 (2). *Ctbp1* and its paralog *Ctbp2* are expressed in overlapping regions along the anterior-posterior axis during development (2). *Ctbp2* is critical for heart development, and *Ctbp2*^*null/null*^ mouse mutants die by mid-gestation at embryonic (E) 10.5 (2). Consistent with their overlapping expression patterns, *Ctbp1* ^*null/null*^; *Ctbp2* ^*null/wt*^ embryos die between E15.5-E18.5 and double null *Ctbp1* ^*null/null*^; *Ctbp2* ^*null/null*^ embryos were arrested at the head fold stage with a lack of progression in heart morphogenesis (2).

*Drosophila Ctbp* (*CtBP*) has been shown to be involved in *wingless* (*Wg-Wnt ligand in Drosophila*) signaling and was initially considered as a transcriptional co-repressor (3-5). CTBP is shown to repress transcription by physically interacting with several partners such as Zeb (6, 7), histone deacetylases (8), and histone methyltransferases and demethylases (7). In vertebrates, APC, WNT signaling pathway regulator (APC) acts as an adaptor that brings CtBP and Catenin, beta 1 (CTNNB1, β-Catenin) together. This complex sequesters β-Catenin away from its DNA-binding partner (TCF), effectively shutting down the Wnt signal (9). However, further studies showed that CtBP could act as a repressor or activator based on the context and depending on the presence or absence of Wnt ligands to activate signaling (10-12). With WNT3A ligand stimulation, CTBP1 upregulates *β-Catenin* expression in colorectal carcinoma cells (13). Thus, the interaction between CTBP1 and WNT-signaling is dynamic and varies based on the context and status of Wnt signaling.

Mutations in the *CTBP1* gene lead to a very rare autosomal dominant neurodevelopmental disorder known as Hypotonia, Ataxia, Developmental Delay and Tooth Enamel Defect Syndrome (HADDTS, OMIM: 617915) (14-20). Pierre Robin sequence (PRS) is a clinically heterogeneous craniofacial disorder characterized by micrognathia, glossoptosis, and upper airway obstruction, frequently accompanied by cleft palate, and may occur as either an isolated anomaly or part of a broader syndromic condition (21). Despite the identification of numerous genes associated with mandibular development and craniofacial morphogenesis, the genetic basis of many cases remains incompletely understood. We have been doing exome and genome sequencing on a research basis to characterize patients with craniofacial malformations and no fully explanatory genetic diagnosis. We report here a proband exhibiting micrognathia with compound heterozygous inherited missense mutations in the *CTBP1* gene and a heterozygous early terminal mutation in the *alkaline phosphatase, liver* (*ALPL*) gene. We created mouse alleles for the *Ctbp1* variants to test the hypothesis these were involved in micrognathia pathogenesis.

## Materials and Methods

### Human subjects and variant discovery

Genetic analysis was performed under a protocol approved by the CCHMC IRB. Written informed consent/assent was obtained from all participants. Exome data was analyzed using standard protocols and GoldenHelix VarSeq software analysis packages. Variants were filtered based on population frequencies, inheritance patterns, allele frequency and predicted pathogenicity scores.

### Mice and animal husbandry

The null allele of *Alpl* (*Alpl*^*tm1Sor*^) was obtained from the Jackson laboratory (RRID:IMSR_Jax: 002741) and mice were genotyped by PCR genotyping with specific primer sets as follows: oIMR0137: ccgtgcatctgccagtttgagggga, oIMR0138: ctggcacaaaagagttggtaaggcag, and oIMR0139: gatcggaacgtcaattaacgtcaat. The breeding was carried out on the C57BL/6J background. Mice were housed with a 12 h light/12 h dark cycle with food and water ad libitum. For timed mating, noon of the day of the vaginal plug was considered as day 0.5. Mouse euthanasia was performed in a carbon dioxide chamber followed by secondary cervical dislocation. All animal work procedures were performed following recommendations in the Guide for Care and Use of Laboratory Animals by the National Institute of Health and approved by the Institutional Animal Care and Use Committee (IACUC) at Nationwide Children’s Hospital (AR21-00067).

### RNA in situ hybridization

RNA in situ hybridization was carried out using RNAscope probes and standard procedures as described before (22). Embryos were dissected at E12.5 and were fixed in formalin for 16–24 h. The tissue was washed in PBS, then dehydrated and embedded in paraffin and were sectioned on the microtome at 5 µm, placed on SuperFrost slides, then baked at 60°C for 1h. Hybridization and amplification steps were performed using the HybEZ oven (Advanced Cell Diagnostics, Newark, CA, USA, ACD) set at 40°C. Manufacturer’s protocol from RNAscope Multiplex Fluorescent Reagent Kit V2 (323100) was followed, TSA Cyanine 3 Fluorophores (NEL744001KT) at 1:750, and a C1 *Ctbp1* custom probe (Catalog # 1038271-C1). All reagents were purchased from Advanced Cell Diagnostics, ACD).

### Generation of *Ctbp1* alleles

*Ctbp1* mice were generated at the Cincinnati Children’s Transgenic Animal and Genome Editing core (RRID:SCR_022642). Mouse zygotes (C57BL/6N strain) were injected with 200 ng/μl CAS9 protein (IDT and ThermoFisher), and reagents to insert the variants: 100 ng/μl Q148H sgRNA (GACTCGAGTGCCTTCCCGAAGGG), 75 ng/μl single-stranded donor oligo-nucleotide (CGGCAGCATCTGTGGAAGAAACGGCAGACTCCACCCTGTGCCACATCCTGAACCTGTACCGACGAACCACCTGGCTACACCAcGCgtTgCGGGAAGGCACTCGAGTCCAGAGTGTAGAGCAGATC; IDT, Iowa), and for the G238S sgRNA (GGTGGTGGTTGTGCTCATTG AGG) and donor (AATCGAGCGGGCCCTGGGGCTACAGCGCGTGAGCACGCTGCAGGACCTGCTCTTCCACAGTGACTGCGTTACCCTGCAcTGCaGCCTgAAcGAGCACAACCACCACCTCATCAATGACTTTACTG) followed by surgical implantation into pseudo-pregnant female (CD-1 strain) mice. PCR genotyping primers for Q148H: CAGTGGCCAACGCACCTAGTCC, and CGTGTGTTTCGTAGGCATCGCAG; G238S: TTGGCTTCAACGTCCTCTTCTATG, and CTCTGAGGGAAGAGTAGAGGGC. Further genotyping was carried out with restriction digests as the Q148H variant creates an *MluI* recognition site and the G238S changes creates a *BtsIv2* recognition sequence. The mouse *Ctbp1* transcript used for reference is ENSMUST00000402685.1.

### Skeletal preparations

Skeletal preparations were carried out using Alcian Blue and Alizarin Red to visualize cartilage and bone, respectively, as per standard procedures. Briefly, animals were skinned, eviscerated and fixed for 2 days in 95% ethanol. Samples were stained overnight at room temperature in 0.03 % (w/v) Alcian Blue solution (Sigma-Aldrich, A3157) containing 80% ethanol and 20% glacial acetic acid and were destained in 95 % ethanol for 24 h followed by pre-clearing in 1% KOH overnight at room temperature. Skeletons were then stained overnight in 0.005% Alizarin Red solution (Sigma-Aldrich, A5533) containing 1% KOH. Samples were then cleared in 20% glycerol/1% KOH solution for 24 h. Finally, they were transferred to 50 % glycerol/50% ethanol for photography. Skeletal preparations were imaged using a Zeiss Discovery.V12 Stereoscope. Mandible and head lengths were measured using Zeiss Zen 3.8 software (Carl Zeiss Microscopy GmbH, Germany).

### Immunofluorescence

Embryo heads were fixed in 4% PFA at 4°C and passed to 15% and then to 30% sucrose for a day each before the embryo heads were embedded in Optimal Cutting Temperature solution (Sakura) and stored at −80°C. 10 μm sections were obtained on positively charged glass slides with a Leica CM 1860 cryostat. Slides selected for immunohistochemistry were dried, rehydrated with PBS. Antibody retrieval was performed with 0.1 M citrate buffer (pH 6.0) for 15 mins and cooled down for 1.5 h as previously described (23). Primary antibody was anti-phospho-histone H3 (pHH3) (Sigma H 0412; 1:500), and cleaved caspase-3 (CC3) (Cell signaling, 9661; 1:250), and the secondary antibody was anti-rabbit IgG Alexa fluor 555 (Invitrogen, A-21428; 1:500). Images were acquired using Zeiss Apotome and positive cells were quantified using ImageJ (24).

### Histology

For comparison of section planes and histological morphology, 10 μm thick sections were subjected to nuclear fast red staining solution, dehydrated through a graded ethanol series and cleared in histoclear, mounted and imaged using Zeiss Apotome.

### Western blot

Mandible tissues were micro-dissected from the embryonic head at E13.5 and snap frozen until further use. Tissues were lysed in RIPA buffer containing protease inhibitor cocktail (Thermofisher). Lysates were centrifuged at 21,000 g for 20 min at 4°C and the collected supernatant was quantified using a BCA assay kit (ThermoFisher). 20 μg of protein was separated on a 4–20% mini-PROTEAN TGX stain-free gradient gel (Bio-Rad) and blotted onto a PVDF membrane using Trans-Blot Turbo transfer system. The membranes were blocked with 5% skim milk. Primary antibodies used were anti-active β-Catenin rabbit polyclonal (Millipore, 1:1000). Chemiluminescent detection visualization on a Bio-Rad ChemiDoc with HRP-conjugate secondary antibody anti-rabbit/mouse IgG, (Abcam, 1:3000). Total protein levels from the stain free gels were used for the normalization of β-Catenin levels. Quantification was carried out using Image lab software (Bio-Rad).

### Cell counts

Cell count analyses were carried out in a blinded approach with the genotype hidden to alleviate observer bias. All counts were carried out using Image J software (24).

### Statistical Analysis

All statistical analyses were performed using Graph pad prism 10 (GraphPad Software, Inc., Boston, MA, USA). One way ANOVA followed by Tukey’s multiple comparison test was carried out to determine p-values between the groups. All graph plotted are mean ± standard error of mean unless otherwise stated.

## Results

### Variants in *CTBP1* identified by exome sequencing

We have been recruiting patients with congenital craniofacial malformations as part of a project studying the human genetics of syndromic craniofacial malformations. One of these patients is a female with Pierre-Robin sequence, cleft palate and plagiocephaly born to unaffected parents. The patient also exhibited a lack of rib mineralization, pulmonary hypoplasia requiring tracheostomy, generalized skeletal dysplasia (osteoporosis, bell-shaped thorax, cervical spine defects), ventriculomegaly with delayed myelination, temporal bone anomalies, and left renal cyst. Here, we focused on identifying the genetic basis for the Pierre-Robin sequence presentation in the patient. Research re-analysis of trio whole exome sequencing was performed and highlighted two different variants in *CTBP1*. The first is a maternal missense allele: chr4:1,216,041:C>T (GRCh38); NM_001012614.2, ENST00000382952.8; c.679G>A; NP_001012632.1:p.Gly227Ser and the other is a paternal missense allele: chr4:1,225,463:C>G (GRCh38); NM_001012614.2, ENST00000382952.8; c.411G>C; NP_001012632.1:p.Gln137His (Figure 1A-D).

**Figure 1.**
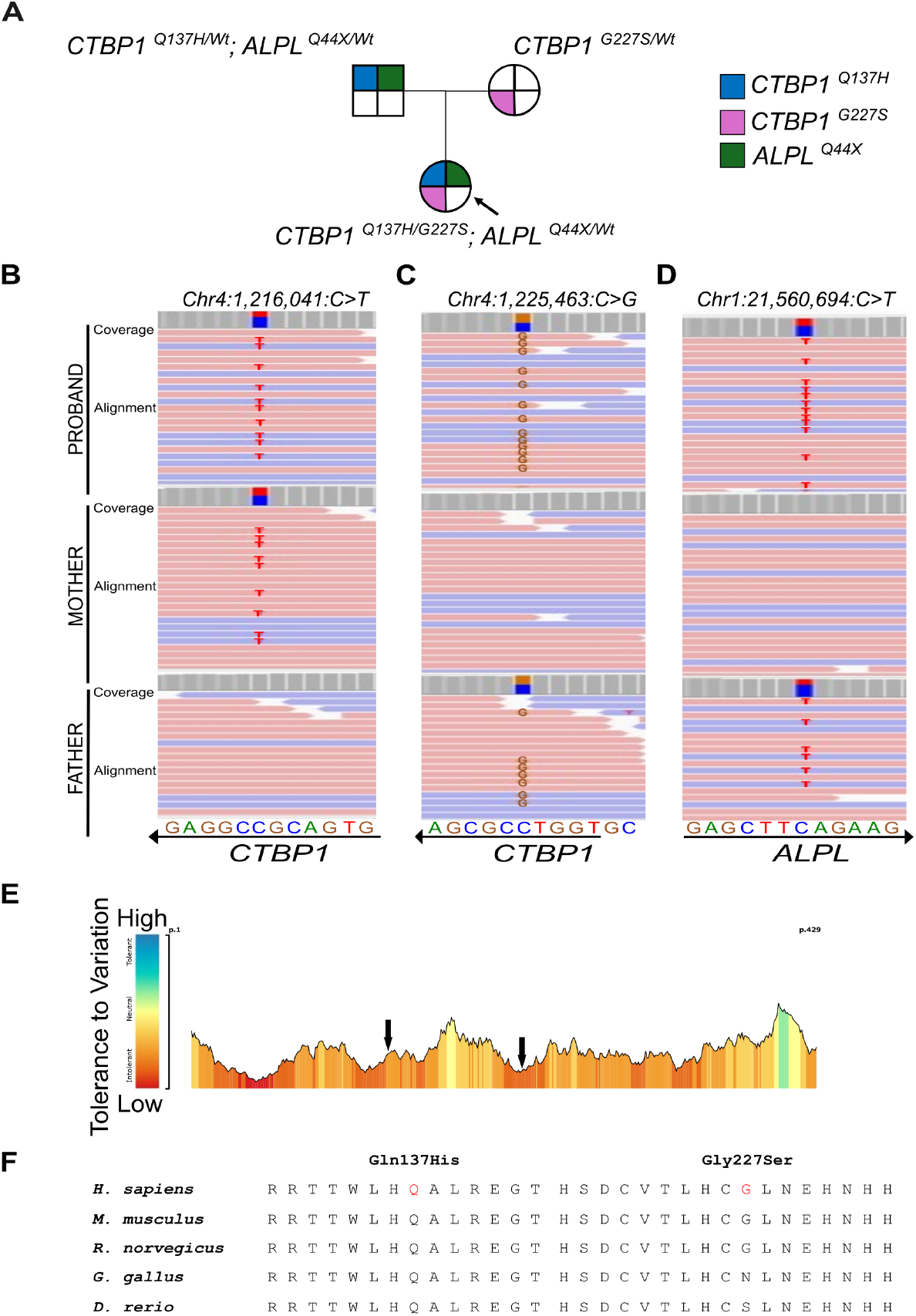
*CTBP1* and *ALPL* variants. (A) Pedigree of sequenced human participants with the affected female proband. Blue colored box denotes *CTBP1*^*Q137H*^, pink denotes *CTBP1*^*G227S*^ and green denotes the *Alpl*^*Q44X*^ variant. Black arrow points to the proband genotype. (B-D) IGV traces highlighting the sequence variants of interest in *CTBP1* (B, C) and *ALPL* (D). (E) MetaDome plot of CTBP1 indicating protein tolerance for variants, black arrows highlight positions of variants identified. (F) Conservation of CTBP1 amino acid sequence around variants of interest.

The Q137H variant is noted in dbSNP (rs1359438837) but is very rare in control populations with an allele frequency of 1.3 × 10^−6^ in gnomAD v4.1.1 (25) (2 alleles of 1,543,164 and no homozygotes), an allele frequency of 1.1 × 10^−5^ in the Regeneron million exome collection (26), 1.4 × 10^−5^ in the NIH All of Us cohort with no homozygotes, and is not noted in the Gabriella Miller Kids First Pediatric Research Program (kidsfirstdrc.org), which has sequence data from hundreds of children with orofacial clefting from multiple ethnicities.

The G227S variant is noted in dbSNP (rs750734000) but is similarly rare in control populations with an allele frequency of 1.5 × 10^−4^ in gnomAD v4.1.1 (25) (245 alleles of 1,611,566 and no homozygotes), an allele frequency of 1.3 × 10^−4^ in the Regeneron million exome collection (26), 8.7 × 10^−5^ in the NIH All of Us cohort with no homozygotes, and 8.4 × 10^−5^ in the Kids First data with 2 of 23,939 alleles and no homozygotes).

The Q137H variant is predicted to be “damaging” by SIFT (27) (score=0.009), CADD (28) (score=24.4) and REVEL (29) (score=0.538) and “damaging” by PolyPhen-2 (30) (score=0.794), and “intolerant” by MetaDome (31) (score=0.42) (Figure 1E). The G227S variant is slightly less pathogenic according to these tools and is predicted to be “damaging” by CADD (score=20.8), “tolerated” by REVEL (score=0.304) and SIFT (score=0.534), “benign” by PolyPhen-2 (score=0.001), and “intolerant” by MetaDome (score=0.20) (Figure 1E). Both amino acids Q137 and G227 are highly conserved among mammals, with amino acid Q137 being totally conserved across vertebrates (Figure 1F). The amino acid change from glycine to serine at position 227 results in altered polarity from a non-polar, hydrophobic residue to polar and hydrophilic and the Missense3D database (32) using the UniProt canonical protein sequence (Q13363, note this is at position 238 in the sequence used for this modeling) predicted the G227S variant to be damaging with the substitution leading to the expansion of cavity volume by 95.688 Å^3 (Figure S1).

Our research analysis also confirmed a variant found in the clinical exome. An early stop mutation in the *ALPL* gene at amino acid 44: Chr1:21560694:C>T (GRCh38); NM_000478.6:c.130C>T:p.(Gln44Ter) was found in both the father and the affected proband. This *ALPL* variant is deemed pathogenic in ClinVar (Variation ID: 370178). Thus, the proband inherited both Q137H and G227S mutations in *CTBP1* from her parents as well as the *ALPL* early stop allele from the father (Figure 1A, D).

Given the craniofacial phenotype of the proband, we examined the expression of *Ctbp1* during mouse embryonic development using RNAscope probes and RNA in situ hybridization at E12.5. *Ctbp1* mRNA was strongly expressed in the mandible and tongue. We also noticed *Ctbp1* expression in the developing brain regions, and optic cup (Figure 2A-C).

**Figure 2.**
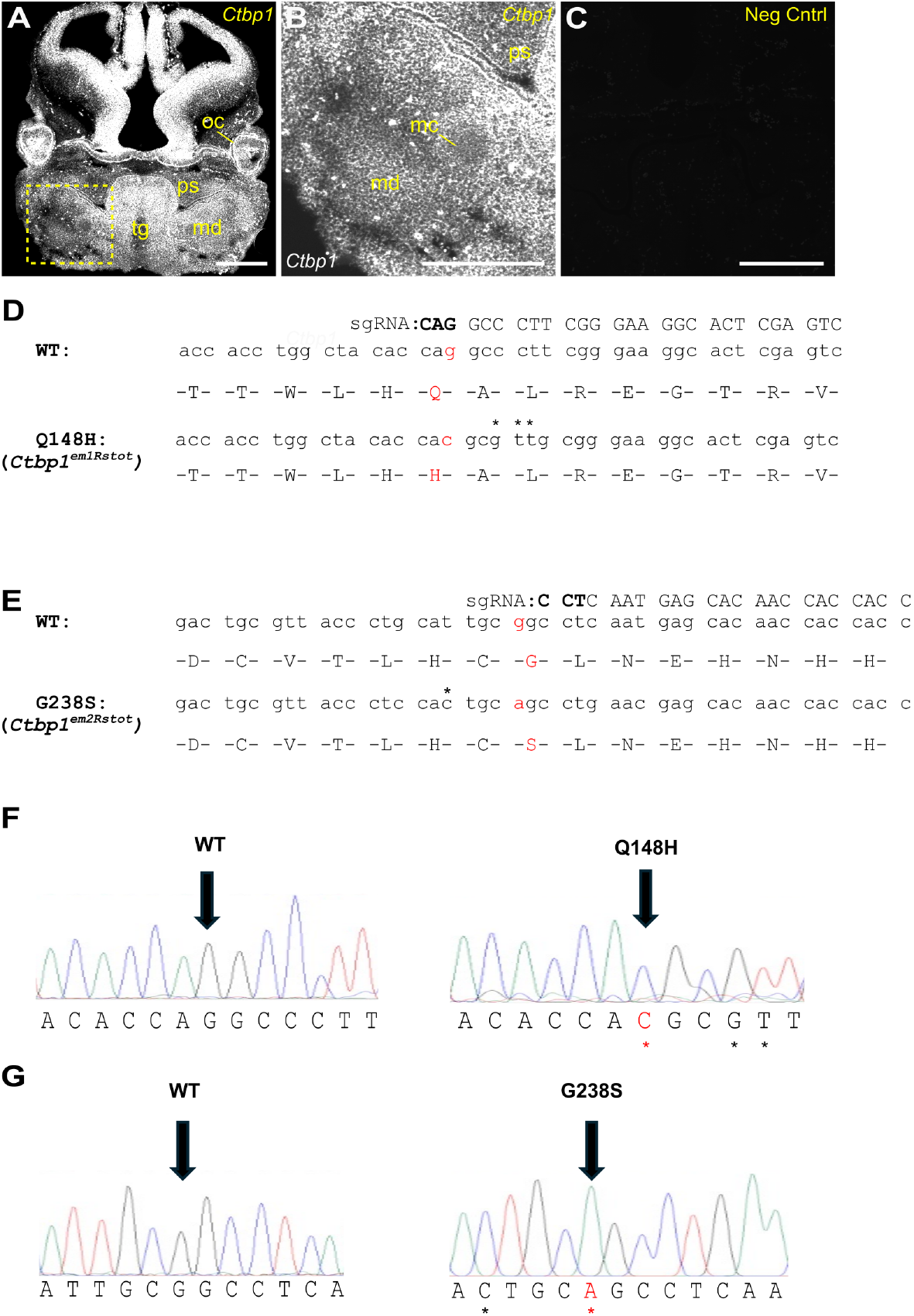
*Ctbp1* expression and generation of mouse alleles. (A–C) RNAscope in situ hybridization showing endogenous *Ctbp1* expression in E12.5 wildtype mouse embryos. *Ctbp1* expression (white) is detected in the developing brain, optic cup, mandible, palatal shelves, and tongue (A, B). The region outlined in yellow in (A) is shown at higher magnification in (B). A negative control probe is shown in (C). mc, Meckels’ cartilage; md, mandible; oc, optic cup; ps, palatal shelves; tg, tongue. Scale bar = 500 μm. (D–G) DNA and amino acid sequences of CRISPR/Cas9-engineered *Ctbp1* alleles and corresponding Sanger sequencing validation traces (F, G). sgRNA target sequences are indicated above the DNA sequence and PAM sites are shown in bold. Black arrows in (F) and (G) indicate the variant nucleotide and the corresponding wildtype nucleotide. Red asterisks denote the engineered missense variants, whereas black asterisks denote silent nucleotide substitutions introduced for genotyping.

### Generation of *Ctbp1* variants in mouse

We used CRISPR/CAS9 genome editing in mouse blastocysts to create two orthologous missense variants in the *Ctbp1* mouse locus corresponding to the variants identified in the proband and validated the targeting by Sanger sequencing (Figure 2D-G). These alleles are *Ctbp1*^*em1Rstot*^ and *Ctbp1*^*em2Rstot*^, hereafter referred to as Q148H, and G238S, respectively (Figure 2D, E). Next, we intercrossed the *Ctbp1*^*Q148H/Wt*^ heterozygous mice and observed that the heterozygous *Ctbp1*^*Q148H/Wt*^ and homozygous *Ctbp1*^*Q148H/Q148H*^ are represented proportionally as expected at weaning (Figure 3A,C; n=55; p=0.741). Similarly, intercrosses of *Ctbp1*^*G238S/Wt*^ mice yielded heterozygous and homozygous offspring at frequencies that did not significantly deviate from expected Mendelian rations, although homozygous *Ctbp1*^*G238S/G238S*^ were recovered at a modestly lower frequency (Figure 3B,D; n=46; p=0.077). To assess the effects of the two *Ctbp1* variants in combination, *Ctbp1*^*Q148H/Wt*^ mice were crossed with *Ctbp1*^*G238S/Wt*^ mice. *Ctbp1*^*Q148H/G238S*^ compound heterozygous offspring were recovered at the expected frequency at weaning (Figure 3E; n=83; p=0.124;). To mimic the proband’s genotype, we incorporated the *alkaline phosphatase, liver/bone/kidney* (*Alpl*) loss-of-function *Alpl*^*null/Wt*^ (*Alpl*^*tm1Sor*^) (33) allele and crossed the *Ctbp1*^*Q148H/Wt*^; *Alpl*^*null/Wt*^ male mice to *Ctbp1*^*G238S/Wt*^; *Alpl*^*Wt/Wt*^ female mice. All expected genotypes were recovered at weaning, although *Ctbp1*^*Q148H/G238S*^; *Alpl*^*null/Wt*^and *Ctbp1*^*G238S/Wt*^; *Alpl*^*null/Wt*^ offspring were observed at modestly lower frequencies than expected (Figure 3F; n=81; p=0.202). Together, these data demonstrate that all modeled allelic combinations are compatible with postnatal survival, enabling further investigation of their effects on craniofacial development (Table 2).

**Table 1.**
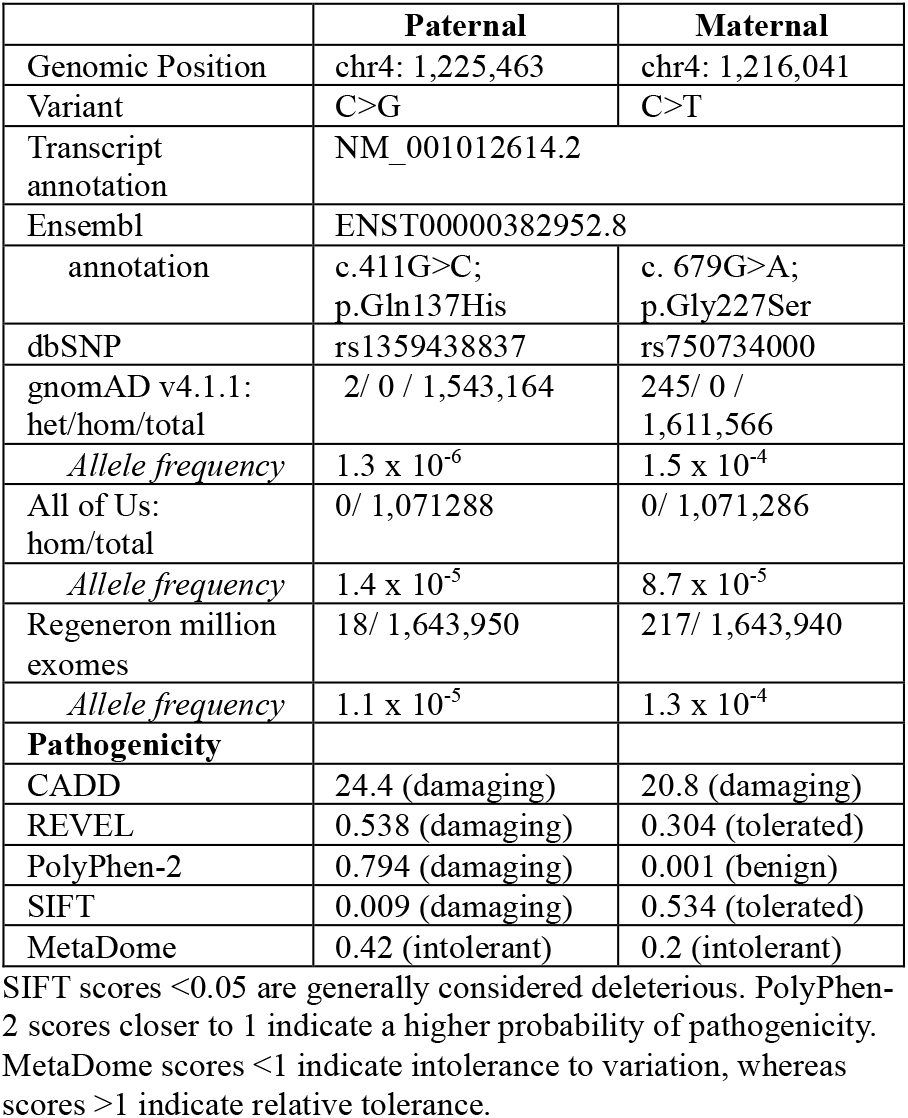
Transcript-specific annotations, population frequencies, and in silico pathogenicity predictions for *CTBP1* variants identified in the proband.

**Table 2.** *Ctbp1, Alpl* mouse survival data.

|  | Age | Total mice | *Ctbp1*<sup>Wt/Wt</sup> | *Ctbp1*<sup>Q148H/Wt</sup> | *Ctbp1*<sup>Q148H/Q148H</sup> |  | p-value<sup>a</sup> |
| --- | --- | --- | --- | --- | --- | --- | --- |
| *Ctbp1*<sup>Q148H/Wt</sup> x *Ctbp1*<sup>Q148H/Wt</sup> | P21 | 55 | 16 | 27 | 12 |  | 0.741 |
|  |  |  | *Ctbp1*<sup>Wt/Wt</sup> | *Ctbp1*<sup>G238S/Wt</sup> | *Ctbp1*<sup>G238S/G238S</sup> |  |  |
| *Ctbp1*<sup>G238S/Wt</sup> x *Ctbp1*<sup>G238S/Wt</sup> | P21 | 46 | 18 | 20 | 8 |  | 0.077 |
|  |  |  | *Ctbp1*<sup>Wt/Wt</sup> | *Ctbp1*<sup>Q148H/Wt</sup> | *Ctbp1*<sup>G238S/Wt</sup> | *Ctbp1*<sup>Q148H/G238S</sup> |  |
| *Ctbp1*<sup>Q148H/Wt</sup> x *Ctbp1*<sup>G238S/Wt</sup> | P21 | 92 | 18 | 29 | 17 | 28 | 0.124 |
|  |  |  | *Ctbp1*<sup>Wt/Wt</sup>; *Alpl*<sup>Wt/Wt</sup> | *Ctbp1*<sup>Wt/Wt</sup>; *Alpl*<sup>null/Wt</sup> | *Ctbp1*<sup>Q148H/Wt</sup>; *Alpl*<sup>Wt/Wt</sup> | *Ctbp1*<sup>G238S/Wt</sup>; *Alpl*<sup>Wt/Wt</sup> |  |
| *Ctbp1*<sup>Q148H/Wt</sup>; *Alpl*<sup>null/Wt</sup> x *Ctbp1*<sup>G238S/Wt</sup> | P21 | 81 | 14 | 8 | 17 | 11 | 0.202 |
|  |  |  | *Ctbp1*<sup>Q148H/Wt</sup>; *Alpl*<sup>null/Wt</sup> | *Ctbp1*<sup>Q148H/G238S</sup>; *Alpl*<sup>Wt/Wt</sup> | *Ctbp1*<sup>G238S/Wt</sup>; *Alpl*<sup>null/Wt</sup> | *Ctbp1*<sup>Q148H/G238S</sup>; *Alpl*<sup>null/Wt</sup> |  |
|  |  |  | 10 | 6 | 7 | 8 |  |

<sup>a</sup>p-values are the result of a Chi-square analysis for each cross

**Figure 3.**
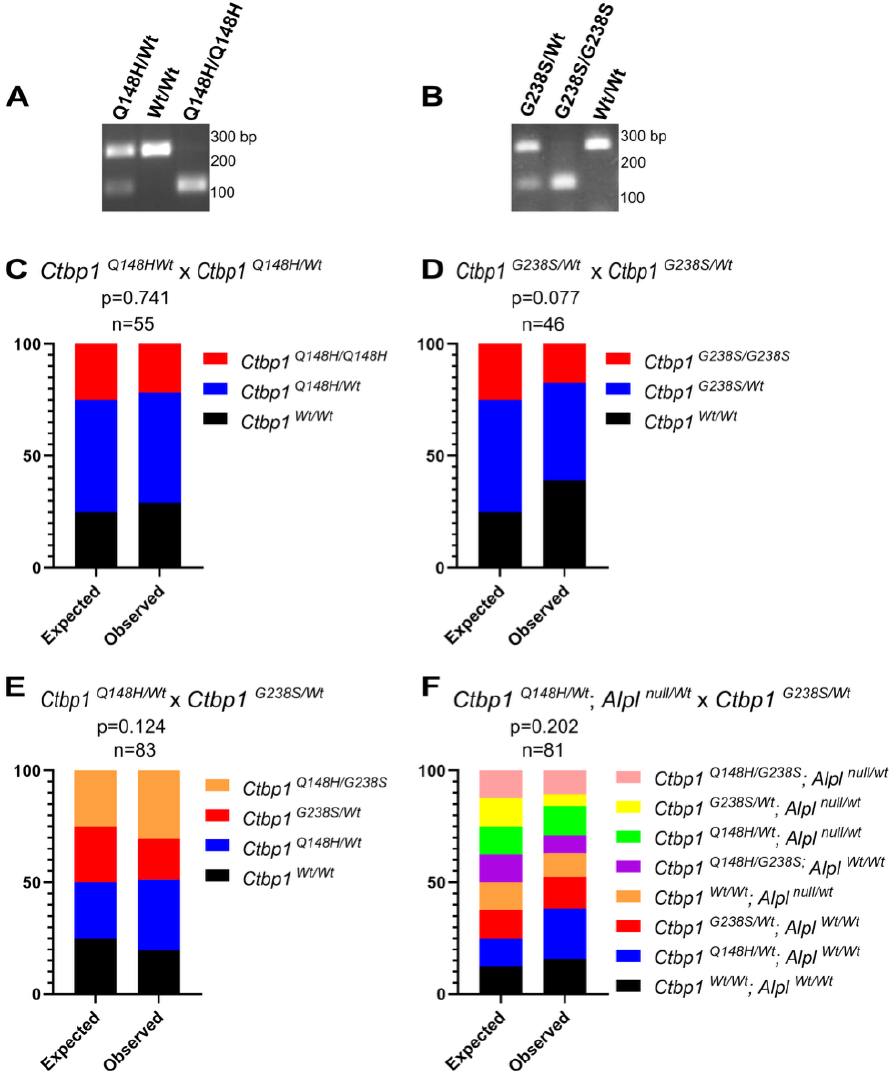
Genotyping and survival of *Ctbp1* alleles at weaning stage. (A, B) Genotyping PCR after *MluI* and *BtsIV2* digestion to genotype for *Ctbp1*^*Q148H*^ (A) and *Ctbp1*^*G238S*^ (B) alleles, respectively. (C-F) Genotypes recovered from crosses of *Ctbp1*(C, D, E) and *Ctbp1; Alpl* (F) alleles indicated at weaning stage. Chi-square p-values are shown for each cross.

### Ctbp1^Q148H/G238S^; Alpl^null/Wt^ mice exhibit mandible defects

Along with kidney and brain abnormalities, the patient also exhibited overt skeletal dysplasia and rib phenotypes. (osteoporosis, bell-shaped thorax, cervical spine defects), ventriculomegaly with delayed myelination, temporal bone anomalies, and left renal cyst. However, gross examination of E18.5 skeletal preparations did not reveal such skeletal dysplasia in the *Ctbp1; Alpl* mutant mouse embryos. We therefore focused our analysis on the mandibular phenotype and examined E18.5 skeletal preparations from embryos carrying various combinations of the *Ctbp1* alleles and the *Alpl*^*null/Wt*^ allele (Figure 4A–D). We quantified total mandibular length, distal mandibular length, proximal mandibular length, mandibular width at two independent positions, and condylar width (Figure 4E). To account for variation in overall head size, mandibular measurements were normalized to head length, as previously described (34). The parental genotypes of *Ctbp1*^*Q148H/Wt*^; *Alpl*^*null/Wt*^ (n=6; p>0.999) and *Ctbp1*^*G238S/Wt*^ (n=4; p=0.149) did not exhibit significant differences in mandibular dimensions compared with wildtype controls (Figure 4F-K). Similarly, *Ctbp1*^*Q148H/G238S*^ compound heterozygous embryos did not show significant reductions in mandibular measurements relative to wild-type littermates (Fig. 4F-K). In contrast, the combination of the three alleles (*Ctbp1*^*Q148H/G238S*^; *Alpl*^*null/Wt*^) resulted in significant reductions in total mandibular length (n = 12; p < 0.001), mandibular width at two independent positions (n = 12; p < 0.001 and p = 0.043), and condylar width (n = 12; p = 0.002) (Figure 4F-K). A trend towards reduced distal mandibular length (n=12; p=0.128) and proximal mandibular length (n = 12; p = 0.056) was also observed in these embryos (Figure 4G,H). Furthermore, we tested the effects of other allelic combinations and found that homozygous *Ctbp1*^*Q148H/Q148H*^; *Alpl*^*null/Wt*^and *Ctbp1*^*G238S/G238S*^; *Alpl*^*null/Wt*^ embryos also exhibited reductions across multiple mandibular measurements. Collectively, these findings support a genetic interaction between *Ctbp1* and *Alpl* and are consistent with a dosage-dependent effect of *Ctbp1* variation on mandibular growth (Figure 5).

**Figure 4.**
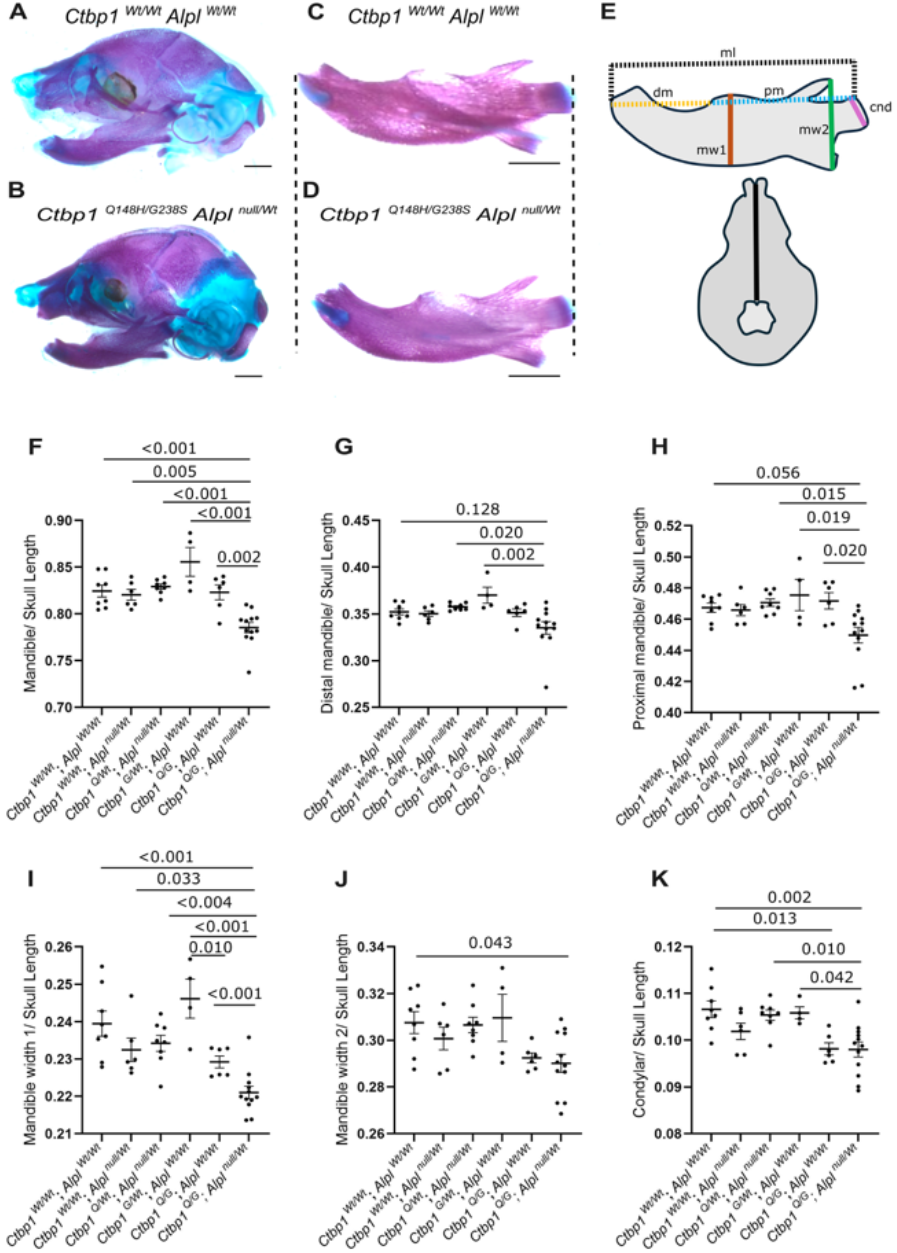
Mandible phenotype in *Ctbp1-Alpl* mutants. (A-D) Skeletal preparations with lateral views of head (A, B) and lingual views of mandible (C, D) in *Ctbp1*^*Wt/Wt*^; *Alpl*^*Wt/Wt*^ (A, C) and *Ctbp1*^*Q148H/G238S*^; *Alpl*^*null/Wt*^ mutants (B, D) at E18.5. Scale bar, 1 mm (E). Schematic showing the different landmarks for the mandible dimension measurements. Black dotted lines show the total mandible length (ml), yellow and blue dashed lines indicate distal mandibular (dm) length and proximal mandibular (pm) length, respectively. The brown and green vertical lines indicate mandibular width 1 (mw1) and mandibular width 2 (mw2), respectively, while the magenta line indicates condylar width (cnd). The lower schematic illustrates the measurement used for normalization of head length. The black vertical line indicates the distance between the distal nasal cartilage and the anterior tip of the basisphenoid bone. ml, total mandibular length; dm, distal mandibular length; pm, proximal mandibular length; mw1, mandibular width 1; mw2, mandibular width 2; cnd, condylar width. (F–K) Quantification of total mandibular length, distal mandibular length, proximal mandibular length, mandibular width 1, mandibular width 2, and condylar width normalized to head length. Data were analyzed by one-way ANOVA followed by Tukey’s multiple-comparison test and are presented as mean ± SEM.

**Figure 5.**
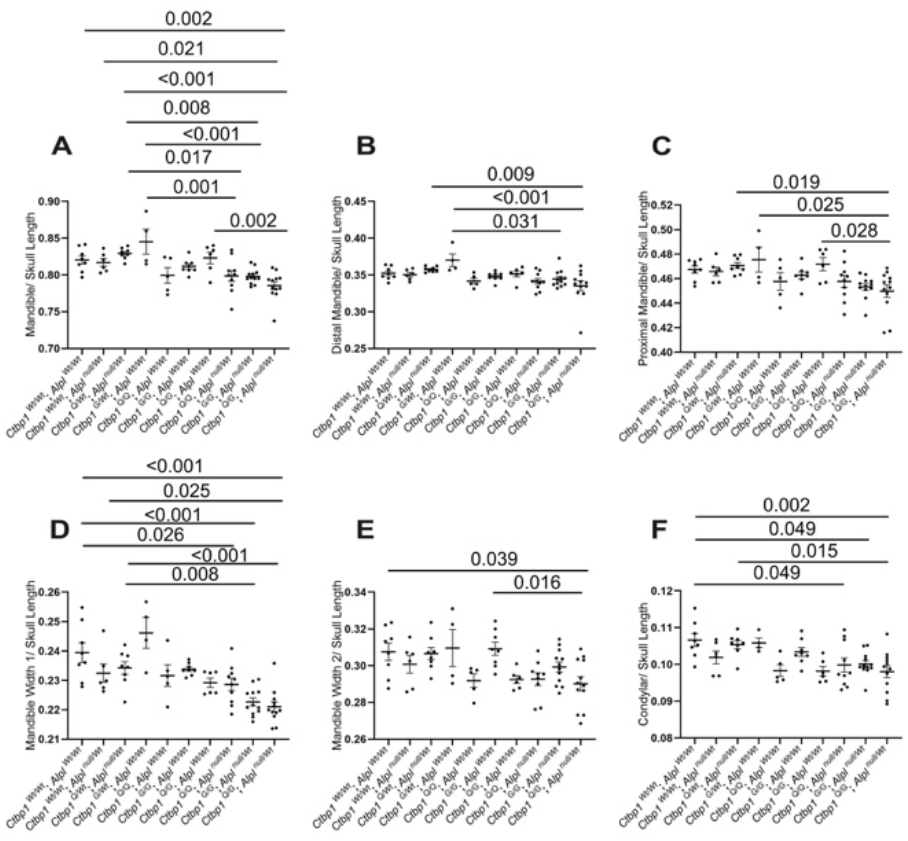
Quantitative mandibular morphometric analyses across *Ctbp1* and *Alpl* allelic combinations. Mandibular dimensions were quantified from E18.5 skeletal preparations and normalized to head length. (A) Total mandibular length, (B) distal mandibular length, (C) proximal mandibular length, (D) mandibular width 1, (E) mandibular width 2, and (F) condylar width in wildtype and mutant embryos carrying the indicated combinations of *Ctbp1* and *Alpl* alleles. Data are presented as mean ± SEM. Statistical analyses were performed using one-way ANOVA followed by Tukey’s multiple-comparison test.

### Effects of *Ctbp1; Alpl* compound alleles on proliferation and cell survival in the developing mandible

To characterize if the mandible growth defect is due to a defect in cell proliferation, we carried out phospho-histone H3 (pHH3) immunostaining in the E13.5 heads of *Ctbp1; Alpl* mutants and counted the pHH3-positive cells in the mandibular region. The relative plane of section is shown in Figure 6M, and representative sections stained with nuclear fast red to illustrate the overall craniofacial morphology and the mandibular region used for quantification are shown in Supplementary Figure S2A, B. While neither parental genotype (*Ctbp1*^*G238S/Wt*^ or *Ctbp1*^*Q148H/Wt*^; *Alpl*^*null/Wt*^) nor the compound heterozygous allele of the *Ctbp1* variants (*Ctbp1*^*Q148H/G238SI*^) exhibited any difference in proliferation, the three-allele combination *Ctbp1*^*Q148H/G238S*^; *Alpl*^*null/Wt*^ alone resulted in a significant reduction in cell proliferation in the mandible (Figure 6A-N; n=4; p<0.001). Analysis of additional genotype combinations further demonstrated that the most pronounced reduction in proliferation occurred in *Ctbp1*^*Q148H/G238S*^; *Alpl*^*null/Wt*^ embryos (Supplementary Figure S3A). To rule out aberrant apoptosis as a cause of the mandible growth deficit, we immunostained for cleaved caspase 3 (CC3) and counted the CC3-positive cells in the mandibular region. Data revealed that the mutant alleles in any combination did not exhibit any increase in apoptosis indicating that the mandible growth defect is unlikely to be due to decreased cell survival (Figure 7A-N; S3B; n=4; p=0.246). Together these data indicate that the variant *Ctbp1*^*Q148H/G238S*^; *Alpl* ^*null/Wt*^ resulted in mandible growth defect with consistent reduction in cell proliferation.

**Figure 6.**
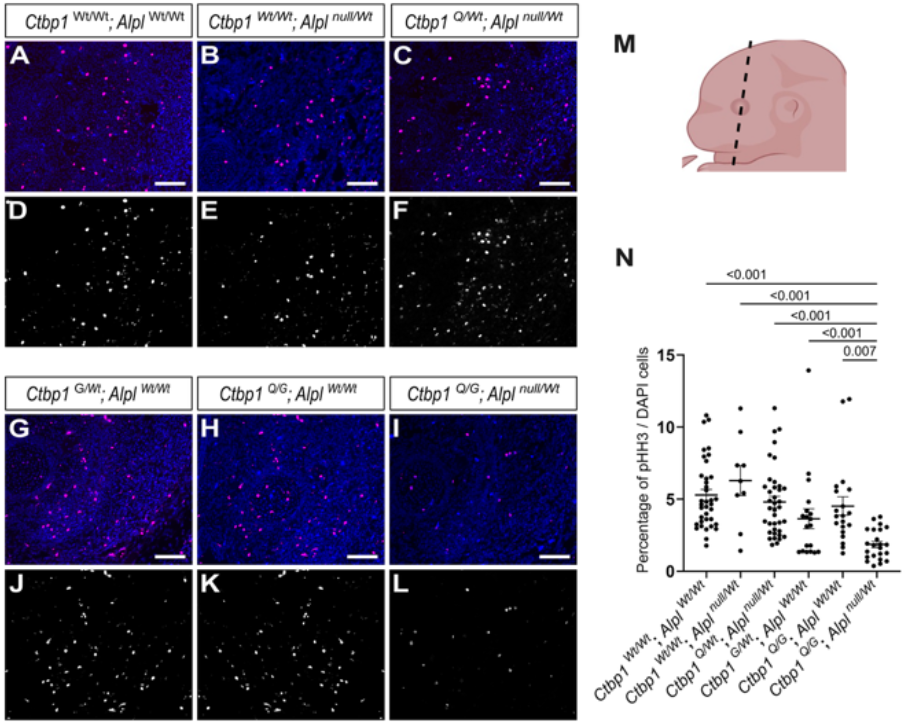
Reduced cell proliferation in the mandibles of *Ctbp1-Alpl* mutants. (A-L) Immunostaining for phospho-histone H3 (pHH3) (A-C, G-I; magenta) overlayed with DAPI (blue) or displayed in white (D-F, J-L) in the coronal sections of mandible at E13.5 in *Ctbp1*^*Wt/Wt*^; *Alpl*^*Wt/Wt*^ (A, D), *Ctbp1*^*Wt/Wt*^; *Alpl*^*null/Wt*^ (B, E), paternal *Ctbp1*^*Q148H/Wt*^; *Alpl*^*null/Wt*^ (C, F), maternal *Ctbp1*^*G238S/Wt*^; *Alpl*^*Wt/Wt*^ (G, J), the compound *Ctbp1*^*Q148H/G238S*^; *Alpl*^*Wt/Wt*^ (H, K), and the proband’s genotype of *Ctbp1*^*Q148H/G238S*^; *Alpl*^*null/Wt*^ (I, L). (M) Schematic of an E13.5 embryo head indicating the plane of section used for analysis (black dashed line). (N) Quantification of pHH3+ cells and data represented as mean ± standard error of mean, one-way ANOVA followed by Tukey’s multiple comparison test was performed for statistical analysis, n=4. Scale bar, 100 μm.

**Figure 7.**
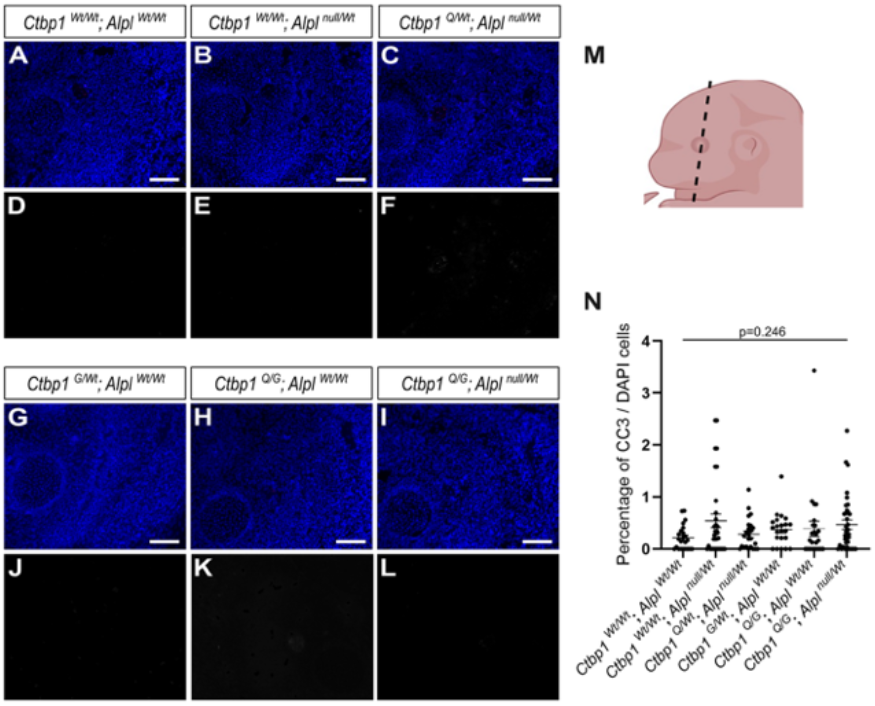
Apoptosis in the mandible of *Ctbp1-Alpl* mutants. (A-L) Immunostaining for cleaved caspase 3 (CC3) (A-C, G-I; magenta) overlayed with DAPI (blue) or displayed in white (D-F, J-L) in the coronal sections of mandible at E13.5 in *Ctbp1*^*Wt/Wt*^; *Alpl*^*Wt/Wt*^ (A, D), *Ctbp1*^*Wt/Wt*^; *Alpl*^*null/Wt*^ (B, E), paternal *Ctbp1*^*Q148H/Wt*^; *Alpl*^*null/Wt*^ (C, F), maternal *Ctbp1*^*G238S/Wt*^; *Alpl*^*Wt/Wt*^ (G, J), *Ctbp1*^*Q148H/G238S*^; *Alpl*^*Wt/Wt*^ (H, K), and the proband’s genotype of *Ctbp1*^*Q148H/G238S*^; *Alpl*^*null/Wt*^ (I, L). (M) Schematic of an E13.5 embryo head indicating the plane of section used for analysis (black dashed line). (N) Quantification of CC3+ cells and data represented as mean ± standard error of mean, one-way ANOVA followed by Tukey’s multiple comparison test was performed for statistical analysis, n=4. Scale bar, 100 μm.

### Ctbp1^Q148H/G238S^; Alpl ^null/Wt^ mutants display reduced β-Catenin levels

*Ctbp1* has been shown to regulate *β-Catenin* and its downstream targets in a context dependent manner (10-12). Moreover, heterozygous *Alpl* (*Alpl*^*null/Wt*^) mice show reduced phospho-GSK3β and active β-Catenin levels in bone marrow stem cells *in vivo*, which were rescued by lithium chloride treatment (35). To assess whether abnormal WNT signaling could be associated with the reduced cell proliferation and mandibular hypoplasia in these mutants, we carried out immunoblotting using an antibody against active CTNNB1 in the E13.5 mandible protein lysates of various combinations of *Ctbp1; Alpl* mutants. We observed a reduction in the active CTNNB1protein levels in the *Ctbp1*^*Q148H/G238S*^; *Alpl*^*null/Wt*^ embryos, while other combinations of the *Ctbp1/Alpl* variants did not exhibit much of a reduction in the active CTNNB1 levels (Figure 8A-C; p=0.026; n=3). These data further show a convergent role for *Ctbp1* and *Alpl* in Wnt/β-Catenin signaling associated proliferation in the mandible during development highlighting the complex genetic nature of these mechanisms involved in the micrognathia phenotype of the proband.

**Figure 8.**
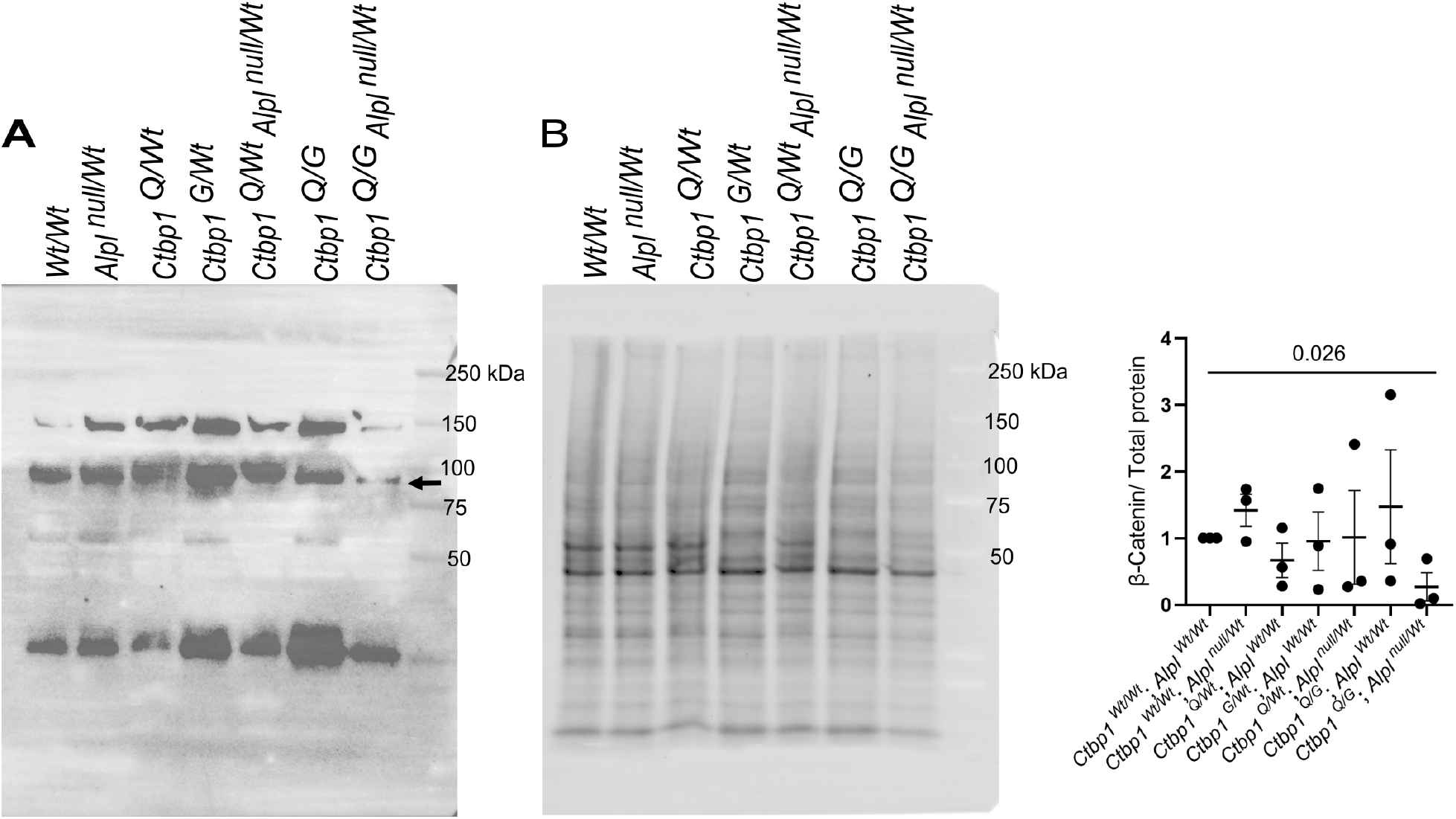
Western blot analysis of active β-Catenin in the mandible of *Ctbp1-Alpl* mutants. (A) Western blot analysis against an active β-Catenin in the mandiblular lysates from the indicated *Ctbp1-Alpl* genotypes at E13.5. Additional bands likely reflect non-specific antibody reactivity or alternative β-catenin isoforms. Only the band corresponding to the expected molecular weight of active β-catenin (arrow) was included in the analysis. (B) The stain-free blots were imaged prior to immunostaining and quantified for normalization. (C) Densitometric analyses of active β-Catenin normalized to total protein and represented as mean ± SEM (n=3).

## Discussion

The widespread adoption of next generation sequencing techniques has resulted in a surge of variants of uncertain significance (VUS). In this study, we have performed trio sequencing on a family where both parents are phenotypically unaffected, but the proband presented with Pierre Robin sequence. The patient carries a complex genotype, inheriting compound heterozygous variants in *CTBP1* (one from each parent) alongside a paternal heterozygous early stop codon in *ALPL*. By generating mouse models for the two *Ctbp1* variants via CRISPR-Cas9 genome editing and combining with an existing null allele of *Alpl*, we have uncovered a plausible mechanism for the micrognathia phenotype observed in the proband. The proband exhibited additional abnormalities, including ventriculomegaly with delayed myelination and skeletal dysplasia. Although overt skeletal dysplasia was not observed in the mutant mice, the present study focused specifically on the mandibular phenotype. We do not see signs of the other patient phenotypes in this mouse model.

The human *CTBP1* variants Q137H and G227S showed varying predictions across multiple in silico tools. Q137H was predicted to be damaging by CADD, REVEL, PolyPhen-2, and SIFT, whereas G227S was predicted to be damaging by CADD but tolerated or benign by REVEL, SIFT, and PolyPhen-2. While MetaDome indicated that both Q137 and G227 are intolerant, the region surrounding Q137 is slightly more tolerant to variation than the region containing G227. However, Q137 itself is highly conserved across vertebrates, and the Q137H substitution is predicted to be damaging by multiple independent prediction algorithms. These observations suggest that regional tolerance estimates may not fully capture the functional consequences of specific amino acid substitutions at highly conserved residues. Missense3D analysis using the UniProt canonical protein sequence (Q13363; corresponding to G238S) further predicted the glycine-to-serine substitution to be damaging, with an expansion of cavity volume. Although G227S is observed at a higher population frequency than Q137H, neither variant has been observed in the homozygous state in the population databases examined.

In the corresponding mouse alleles, homozygous *Ctbp1*^*G238S*^ mice were recovered at modestly lower frequencies than homozygous *Ctbp1*^*Q148H*^ mice. However, both *Ctbp1*^*Q148H/Q148H*^ and *Ctbp1*^*G238S/G238S*^ exhibited hypoplasia of the mandible in combination with the heterozygous *Alpl* allele. Indeed, proliferation is reduced in the oligogenic combination of *Ctbp1*^*Q148H/G238S*^; *Alpl*^*null/Wt*^ mandibles. Thus, while both *Ctbp1*^*Q148H*^ and *Ctbp1*^*G238S*^ in combination with *Alpl* have significant effects in the mandible phenotype, *Ctbp1*^*G238S*^ could have deleterious effects in other systems that affect the survival of the mice.

Several mutations in the *CTBP1* gene lead to a very rare autosomal dominant neurodevelopmental disorder known as HADDTS (14-20). The clinical presentations of our patient are not part of the typical HADDTS observed in other *CTBP1* patients, consistent with a different inheritance pattern. The rarity of the genotype identified in the present study further supports this distinction. Although the individual *CTBP1* variants are present at low frequencies in population databases, neither has been observed in the homozygous state in gnomAD v4. Moreover, we are not aware of additional individuals carrying a comparable combination of *CTBP1* and *ALPL* variants in available sequencing datasets, including the Gabriella Miller Kids First Research Program. Although the *Alpl*^*tm1Sor*^ allele may not fully reproduce the molecular consequences of the human *ALPL* p.Gln44Ter variant, both are expected to result in substantial loss of ALPL function. The *Alpl*^*tm1Sor*^ allele is a well-characterized null allele generated through targeted deletion of exons 2–6 and was selected to model substantial loss of ALPL function *in vivo*. The proband exhibited additional clinical features beyond micrognathia and Pierre Robin sequence, including skeletal, pulmonary, renal, and neurodevelopmental abnormalities. While we did not investigate neurodevelopmental abnormalities, in this study we did not observe any overt skeletal phenotypes or rib dysplasia. Additional studies will be required to determine the extent to which these variants influence other aspects of the proband phenotype.

Definitive oligogenic inheritance for a phenotype is still relatively rare. An early report on Bardet-Biedl syndrome (BBS) with pigmentary retinal dystrophy, polydactyly, obesity, developmental delay, and renal defects associated compound mutations in genes *BBS2* and *BBS6* with three alleles required to manifest the phenotype (36). As more sequencing has been performed and diagnoses made, BBS is canonically now considered a recessive disease with modifying loci (37). Interestingly, oligogenic inheritance of heterozygous variants in three different genes *MKL2, MYH7*, and *NKX2-5* together were shown to cause cardiomyopathy (38). These complex inheritance patterns are rare and make it particularly challenging to delineate the mechanistic interactions and convergent signaling pathways underlying disease phenotypes. However, these oligogenic inheritance patterns may represent a significant portion of currently unexplained cases and providing supporting evidence as we do here is quite resource intensive.

Variants in WNT signaling are robustly associated with micrognathia phenotypes. Autosomal dominant Robinow Syndrome is associated with pathogenic variants in several components of WNT signaling, including missense variants in *WNT5A* and characteristic frameshift variants in *DVL1, DVL3* or *FZD2* (39-43). A pathogenic truncating variant in *CTNNB1* (*β-Catenin*) has also been reported in an individual with micrognathia and neurodevelopmental conditions (44). Both *Ctbp1* and *Alpl* converge on β-Catenin/Wnt signaling. Heterozygous *Alpl* (*Alpl*^*null/Wt*^) mice show reduced phospho-GSK3β and active β-Catenin levels in bone marrow stem cells *in vivo*, which were rescued by lithium chloride treatment (35). Furthermore, CTBP1 has been known to play a context dependent role based on the presence or absence of Wnt ligands to activate signaling (10-12). With WNT3A ligand stimulation, *Ctbp1* induces *β-Catenin* expression (13). Thus, the interaction between *Ctbp1* and Wnt-signaling is dynamic and varies based on the context and presence of ligand. Mice with both variants of *Ctbp1* and the *Alpl* allele showed downregulation of active β-Catenin in the mandible during development. Together these data reveal a complex interplay between *CTBP1* and *ALPL* variants involved in the mandibular hypoplasia with convergence at the level of Wnt/β-Catenin signaling.

## Acknowledgement

We appreciate comments on the manuscript from the Stottmann laboratory group and periodic re-analysis of all sequenced cases in our cohort by I. Showpnil. Funding for this project comes from the National Institutes of Health, R01DE027091, and Abigail Wexner Research Institute at Nationwide Children’s Hospital recruitment funds (R.W.S.). No competing interests are declared.

---

**Supplementary Figure 1.**
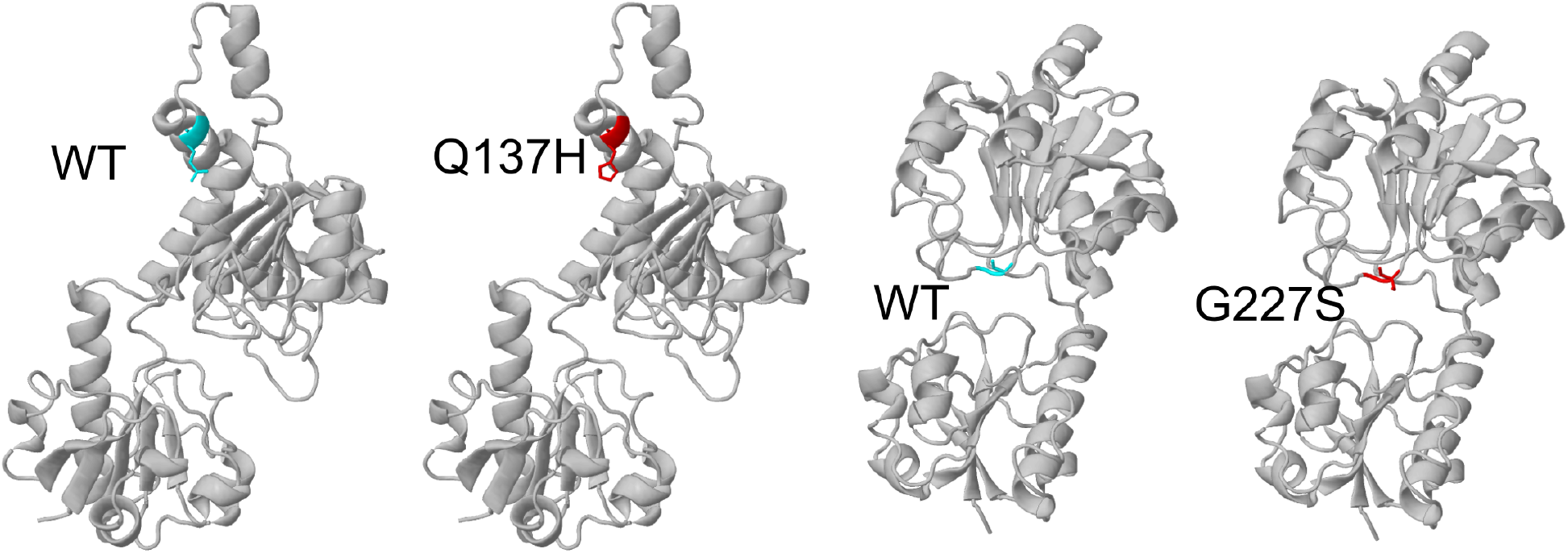
Predicted structural consequences of the CTBP1 Q137H and G227S variants as determined by Missense3D.

**Supplementary Figure 2.**
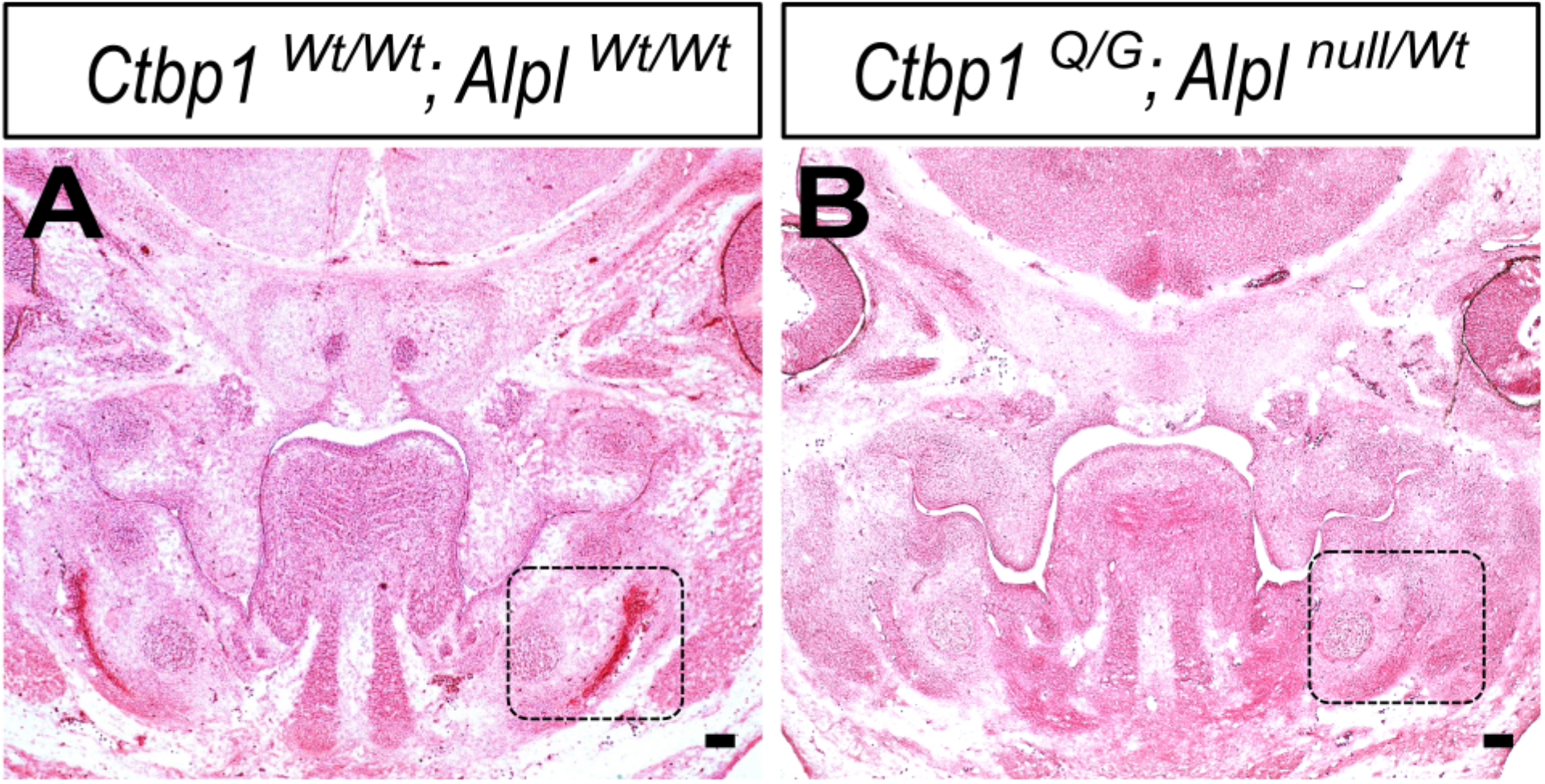
Representative coronal sections through the craniofacial region of *Ctbp1*^*Wt/Wt*^; *Alpl*^*Wt/Wt*^ (A) and *Ctbp1*^*Q148H/G238S*^; *Alpl*^*null/Wt*^ (B) E13.5 embryo heads stained with nuclear fast red. Black dashed boxes in (A) and (B) indicate the mandibular regions used for quantification of pHH3- and CC3-positive cells in Figure 6 and 7, respectively. Scale bar, 100 μm.

**Supplementary Figure 3.**
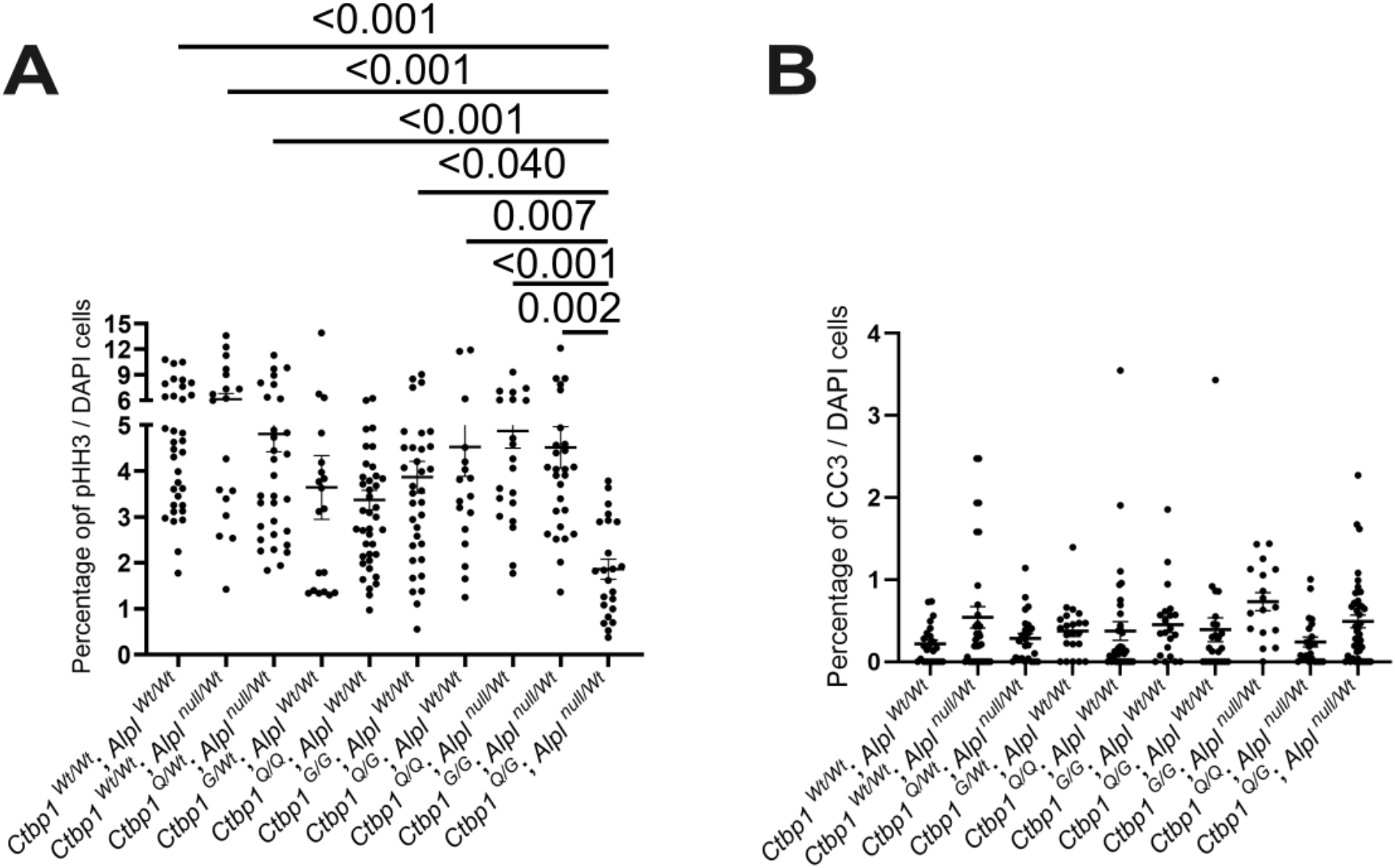
Quantification of pHH3-positive (A) and cleaved caspase-3 (CC3)-positive (B) cells in coronal sections from embryos carrying the indicated *Ctbp1* and *Alpl* genotypes. Data are presented as mean ± SEM. Statistical analyses were performed using one-way ANOVA followed by Tukey’s multiple-comparison test (n = 4).

